# MiRNA expression profiling and zeatin dynamic changes in a new model system of *in vivo* indirect regeneration of tomato

**DOI:** 10.1101/2020.08.03.234013

**Authors:** Huiying Cao, Xinyue Zhang, Yanye Ruan, Lijun Zhang, Zhenhai Cui, Xuxiao Li, Bing Jia

## Abstract

Callus formation and adventitious shoot differentiation could be observed on the cut surface of completely decapitated tomato plants. We propose that this process can be used as a model system to investigate the mechanisms that regulate indirect regeneration of higher plants without the addition of exogenous hormones. This study analyzed the patterns of trans-zeatin and miRNA expression during *in vivo* regeneration of tomato. Analysis of trans-zeatin revealed that the hormone cytokinin played an important role in *in vivo* regeneration of tomato. Among 183 miRNAs and 1168 predicted target genes sequences identified, 93 miRNAs and 505 potential targets were selected based on differential expression levels for further characterization. Expression patterns of six miRNAs, including sly-miR166, sly-miR167, sly-miR396, sly-miR397, novel 156, and novel 128, were further validated by qRT-PCR. We speculate that sly-miR156, sly-miR160, sly-miR166, and sly-miR397 play major roles in callus formation of tomato during *in vivo* regeneration by regulating cytokinin, IAA, and laccase levels. Overall, our microRNA sequence and target analyses of callus formation during *in vivo* regeneration of tomato provide novel insights into the regulation of regeneration in higher plants.

## Introduction

Tissue culture established over 150 years ago continues to play an important role in plant propagation, and continues to be utilized in both basic and applied plant research, including gene transformation and molecular breeding [1, 2]. In-depth studies into mechanisms of regulation of regeneration of higher plants using *in vitro* culture techniques, identified several proteins and transcription factors such as WUSCHEL (WUS), SHOOT MERISTEMLESS (STM), BABY BOOM (BBM), and MONOPTEROS (MP) [3–6]. However, both direct regeneration and indirect regeneration via an intermediate callus phase are introduced by various plant growth regulators supplemented media in traditional tissue culture. Yin[7] reported that 157 unique proteins were significantly differentially expressed during callus differentiation in rice when treated with different relative concentrations of the hormones cytokinin and auxin. Additionally, even though somatic embryogenesis (SEG) has been proposed to be a model system of plant embryogenesis, the expression of gene families such as those of MIR397 and MIR408 was detected in somatic embryos (SE), but greatly decreased in zygotic embryos (ZE) in conifer species[8].

An interesting phenomenon has been observed that *in vivo* adventitious shoots can be regenerated from cut surfaces of stems or hypocotyls after removal of both apical and axillary meristems in some species such as Cucurbita pepo[9], tomato[10], and poinsettia [11]. In tomato, the surface of cut stems regenerates plenty of shoots via callus formation. This *in vivo* generation does not depend on the presence of exogenous hormones. We propose that this phenomenon is particularly useful as a model system to study the innate molecular mechanisms of plant regeneration MicroRNAs (miRNAs) are a class of small noncoding-RNAs (20–24 nt) that regulate gene expression at post-transcriptional levels by directly binding to their targets[12, 13]. In the past 20 years, miRNAs have been shown to play key roles at each major stage of plant development [14–17]. Furthermore, recent studies have shown that miRNAs are involved in callus initiation, formation and differentiation. For example, the expression levels of miR408, miR164, miR397, miR156, miR398, miR168, and miR528 were up-regulated during maize SE induction[18]. Another study demonstrated that over-expression of miR167 inhibited somatic embryo formation by inhibiting the auxin signaling pathway in *Arabidopsis*[19]. In citrus, the ability of the callus to form SEs was significantly enhanced by either over-expression of csi-miR156a or by individual knock-down of its two target genes, *CsSPL3* and *CsSPL14*[20].

Recently, the large scale application of next-generation sequencing has proved to be a useful tool to identify the patterns of miRNA expression during plant regeneration. Genome-wide miRNAs and their targets have been analyzed during explant regeneration in vitro in wheat[21], rice[22, 23], cotton[24], peanut[25], sweet orange [26], coconut[27], larch[28], maritime pine[8], Norway Spruce[29], longan[30], yellow-poplar[31], radish[32], Lilium [33], and Tuxpeno maize[34]. However, all these studies were performed on *in vitro* specimens, which relys on the presence of exogenous hormones to regenerate plantlets. The aim of the present study was to identify the pattern of miRNA expression during callus formation in *in vivo* regeneration in tomato through sequencing. We assessed 92 known miRNAs and identified 91 novel miRNAs, of which several were found to be developmentally regulated. We also analyzed dynamic changes in cytokinin levels during *in vivo* regeneration of tomato.

## Materials and Methods

### Plant materials

The tomato cultivar micro-TOM was used in this study. Seeds were placed on moistened filter papers for approximately 3 to 4 d until the seeds sprouted. The germinated seeds were seeded into a tray of 72 cells filled with a mixture of nutrient soil, matrix, vermiculite and perlite (2:2:1.5:0.5(v/v/v/v)), and grown in a culture room with temperature ranging from 23 to 28 °C and 16/8 h light/dark photoperiod. When the seedlings had grown 6 to 8 true leaves, the primary shoot was decapitated horizontally. All axillary buds that appeared after decapitation were resected at the base.

### HPLC analysis of trans-zeatin

Cutting surfaces of stems (3 mm long) were sampled at 0, 9, 12, 15, 18, 21, 24 and 30 d after decapitation. The samples were immediately frozen in liquid nitrogen and stored at −80 °C. Trans-zeatin was extracted with 80% methanol from samples and analyzed using HPLC (Agilent 1100) connected to an UV detector (λ = 274 nm). The passing fraction was further purified by Sep-pak C18 column (Waters). Gradient elution was with a mixture of water-methanol (75:25 (vol:vol)) with an elution rate of 1.0 mL/ min at a column temperature of 35 °C. The absorbing material was Agilent C18 with a particle size of 5 μm loaded into a stainless steel column (250 × 4.6 mm).

### Lovastatin addition during the tomato regeneration in vivo

Lovastatin (Sigma-Aldrich, USA) dissolved in DMSO (stock solution: 0.124 M) and the final concentrations of 123.6μM was applied. 20μL lovastatin was dripped on cutting surfaces of 10 tomato stems after decapitation with 1μL Tween-20. Another 10 tomato plants were treated with water as control.

A lovastatin (Sigma-Aldrich, USA) stock solution at a concentration of 0.124 M was prepared by dissolving in DMSO and applied at a final concentration of 123.6 μM. A solution of 20 μL lovastatin and 1 μL Tween-20 was applied on the cut surfaces of 10 tomato stems after decapitation. An additional equal number of tomato plants were treated with water as control.

### sRNA library construction and RNA sequencing

Total RNA was extracted from decapitated stem at 0 and 15 d in three biological replicates. Each biological replicate was from a pool of 8–10 tomato plants. RNA samples of the three biological replicates were mixed in equal amount and used for the construction of libraries. The small RNA library was constructed using 3 μg total RNA from each treatment respectively as input materials. The sequencing library was generated by NEBNext^®^ Multiplex Small RNA Library Prep Set for Illumina^®^ (NEB, USA), with added index codes to attribute sequences of each sample as recommended by the manufacturer. Briefly, the NEB 3′ SR Adaptor was connected to 3′ end of miRNA, siRNA and piRNA directly. After the 3′ ligation reaction, the single-stranded DNA adaptor was transformed into a double-stranded DNA molecule by hybridization of the SR RT Primer with excess of 3′ SR Adaptor (kept free after the 3′ ligation reaction). This step significantly reduced the formation of adaptor-dimers. In addition; dsDNAs were not the T4 RNA Ligase 1-mediated-substrates, and therefore were not ligated to the 5′ SR Adaptor in the following ligation step. The 5′ ends adapter was connected to the 5′ ends of miRNAs, siRNA, and piRNA. Reverse transcription reaction was performed using M-MuLV Reverse Transcriptase (RNase H^−^) after ligation with adapters, and Long Amp Taq 2X Master Mix, SR Primer for Illumina and index (X) primer was used for PCR amplification. An 8% polyacrylamide gel was used for purifying PCR products, small RNA fragments approximately 140–160 bp were recovered and dissolved in elution buffer. Finally, the quality of library was evaluated on the 2100 system of Agilent Bioanalyzer using DNA High Sensitivity Chips. TruSeq SR Cluster Kit v3-cBot-HS (Illumina, San Diego, CA, USA) was used to evaluate index-coded samples on a cBot Cluster Generation System. After clustering, the library preparations were sequenced on an Illumina Hiseq 2500 platform, and 50 bp single-end reads were generated.

### MiRNA identification and target prediction

All sequenced data were firstly filtered with the removal of N% >10% reads, length <18 nt or >30 nt, with 5′ adapter contamination, 3′ adapter null or insert null and low quality reads to obtain clean reads. The remaining clean reads were mapped to the reference sequence by Bowtie to obtain unique reads and analyze the length distribution and expression of unique sRNAs. Unique reads were mapped to miRNA, rRNA, tRNA, snRNA, snoRNA, repeat masker, NAT, TAS, exon, intron, and others. Mapped sRNA tags were used to search for known miRNAs. With miRBase20.0 as reference, the potential miRNAs were identified using modified software mirdeep2 and srna-tools-cli, and then the secondary structures were drawn. We used miREvo and mirdeep2 to predict novel miRNAs through the precursor structure of each miRNA unannotated in the previous steps, including the analysis of the secondary structure, the dicer cleavage site, and the minimum free energy. We used miFam.dat (http://www.mirbase.org/ftp.shtml) to compare our candidate miRNA families with known miRNA families from other species.

We used psRobot_tar in psRobot to identify the potential gene targets of known and novel miRNAs.

### Differential expression analysis of miRNAs

We used the TPM (transcript per million) value to estimate the differential expression levels of miRNAs between stem and callus[35]. The TPM ratio of miRNAs between stem and callus libraries was computed as log_2_ (callus/stem). miRNAs with p value <0.05 and log_2_ (callus/stem) <−1 or >1 were regarded to have as significantly differential expression levels through the Bayesian method.

### MiRNAs quantification by qRT-PCR

Primers were designed by stem-loop method to perform RT-PCR assays according to Chen’s design[36]. A 20 μL final volume Reverse transcription (RT) reaction was carried out to validate the expression levels of selected miRNAs extracted from stem and callus respectively. The final 20 μL reaction system included 1 μL template, 1 μL stem-loop primer, 1 μL dNTP mixture, 4 μL buffer, 1 μL reverse transcriptase and 0.5 μL RNase inhibitor. In the reaction tube, stem-loop primer, dNTP mix and template were added first, and then the template was denatured at 65 °C for 5 min to improve the efficiency of reverse transcription. The tube was placed on ice to cool for 2 min, followed by the addition of the buffer, reverse transcriptase and RNase inhibitor. The tube was then incubated at 45 °C for 60 min, followed by 95 °C for 5 min. The reverse transcription reaction was completed by cooling on ice for 2 min. After reverse transcription, 1 μL of the RT reaction mixture was used for PCR. The PCR system was 25 μL, containing 12.5 μL PCR mix, 1 μL template, 1 μL downstream primer and 1 μL upstream primer, supplemented to 25 μL with nuclease-free water. The PCR conditions were as follows: 94 °C for 2 min, 94 °C for 30 s, 60 °C for 1 min for 35 cycles, followed by a final extension of 72 °C for 5 min. Following the PCR assay, gel electrophoresis was used to detect the amplified products.

Quantitative reverse transcription-polymerase chain reaction (qRT-PCR) assays were performed using a C1000 Touch^TM^ Thermal Cycler (Bio-Rad, Hercules, CA, USA). The reaction system included 5 μL SYBR^®^ Premix Ex Taq^TM^ (Takara, China), 1 μL template, 0.2 μL upstream primer, 0.2 μL downstream primer and 3.6 μL nuclease-free water. The reaction conditions were as follows: 94 °C for 2 min, 94 °C for 30 s, 60 °C for 1 min, and final extension at 72 °C for 5 min for 35 cycles. The sequences of all primers used in this study are compiled in S1 Table.

## Results

### Phenotypic analysis of in vivo regeneration

In tomato, a primary shoot shows apical dominance and inhibits outgrowth of axillary buds. After excising the main shoot apex, the dormant axillary buds began to develop immediately to replace the lost shoot apex. Since all new axillary buds were excised, light-green callus gradually formed at the cut surface of primary shoots and axillary buds followed by progression to the compact and nodular stage with a maximum diameter of up to 1 cm. When the callus entered its differentiation stage, a large number of purple dots appeared on its surface, and finally the shoots appeared to regenerate through callus (Fig. 1). It took 30 days to obtain macroscopic shoots after decapitation at 25°C. These features of *in vivo* regeneration were similar to the responses seen in tissue culture (Tezuka et al., 2011).

**Fig. 1.**
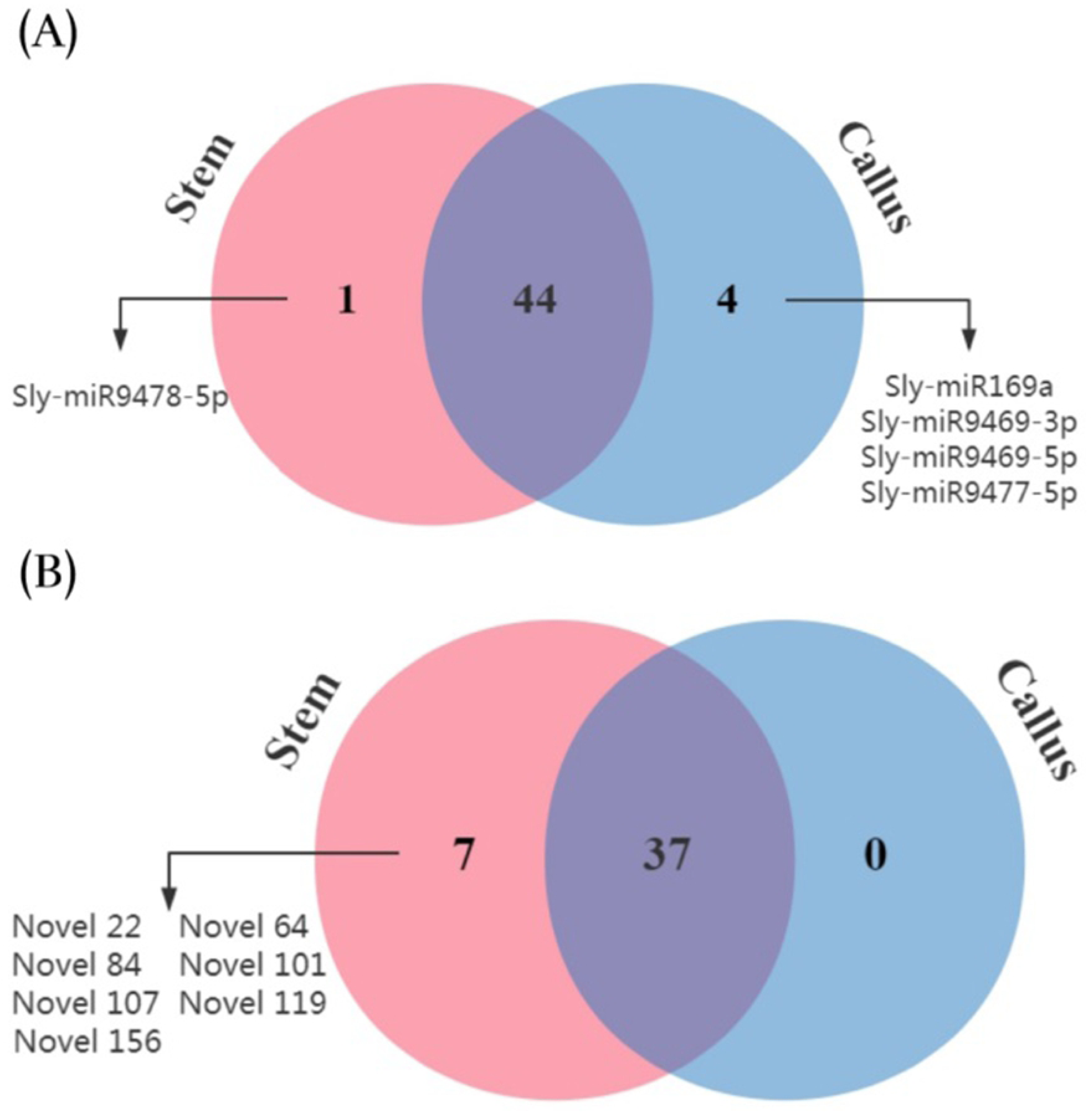
External appearance of the different stages of *in vivo* regeneration in tomato. (A) The decapitated primary shoot; (B),(C) The callus formed on the cutting surface at 15 and 25d after decapitation; (D) The adventitious shoots differentiated from callus.

### Analysis of trans-zeatin during in vivo regeneration

HPLC analysis of trans-zeatin during *in vivo* regeneration of tomato micro-TOM is presented in Fig. 2. Trans-zeatin was not detected in 0 d stem, but was detected in gradually increasing amounts correlated with the progress of callus initiation, formation and differentiation. These results show that cytokinin plays a key role during *in vivo* regeneration of tomato. These data correlated well with previous studies showing that 6-benzyladenine treatment increased the number of adventitious shoot amounts during *in vivo* tomato regeneration [37]. Furthermore, our findings are also supported by other studies which used zeatin as the only exogenous hormone during *in vitro* regeneration of tomato [38–40].

**Fig. 2.**
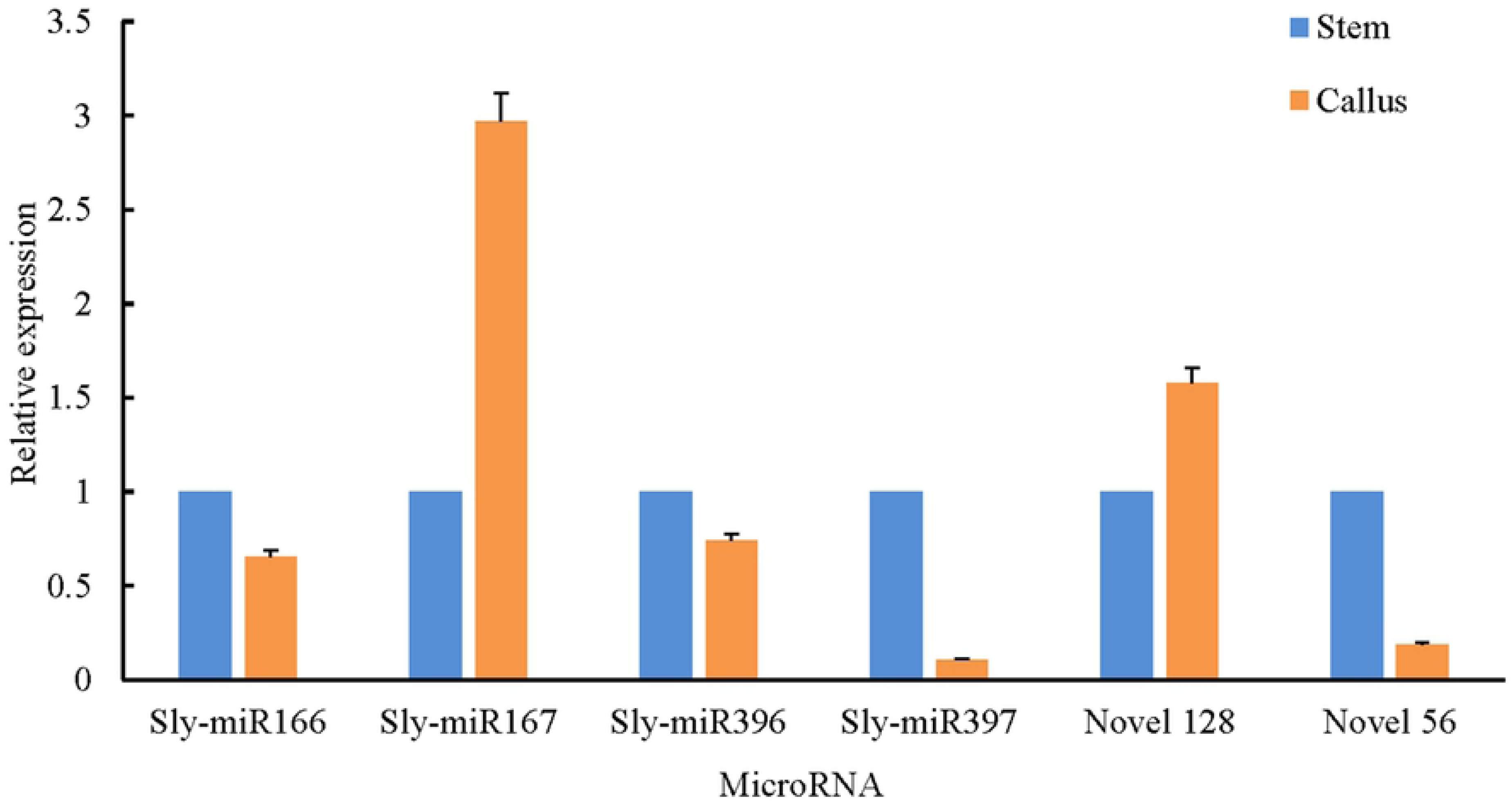
HPLC analysis of trans-zeatin levels during *in vivo* regeneration of tomato micro-TOM. The samples were the cutting surface of stem at 0, 9, 12, 15, 18, 21, 24 and 30 d after decapitation.

Cytokinins are a heterogenous group of N6-substituted adenine derivatives[41]. Lovastatin is a potent inhibitor of the mevalonate pathway, and in principle blocks the synthesis of isopentenyl-pyrophosphate and inhibit the biosynthesis of cytokinin[42]. Lovastatin (1 µM) has been shown to completely inhibit the growth of cultured tobacco cells [43]. However, in this study, the addition of high levels lovastatin (123 µM) to the cut surface of decapitated stems did not inhibit tomato regeneration *in vivo*. There was no obvious difference in the number of regenerated adventitious shoots between lovastatin and control treated plants. Together, these observations suggested that cytokinin was not biosynthesized de novo in the cells at the cut surface or in the callus during *in vivo* regeneration. Cytokinin is found in the xylem sap as a long-distance signal in intact plants [44–46]. To date, the trans-zeatin is the major form of cytokinin in xylem sap [47]. The trans-zeatin is produced mainly in the root, and can then be transported from the root to shoot[48]. Therefore, we speculate that cytokinin was also transported over long distances from the root to callus cells during *in vivo* regeneration of tomato.

### Deep-Sequencing of sRNAs in stem and callus

To study the vital role of miRNAs during *in vivo* regeneration, the cut surfaces of stems at 0 and 15 days after decapitation were used to construct two sRNA libraries. Both of these libraries were sequenced with 13.6 and 11.0 million raw reads obtained from stem and callus libraries, respectively (S2 Table). After removal of the low quality reads as described in the Materials and Methods section, 9.5 and 7.5 million clean sRNAs were obtained from the stem and callus libraries, the stem and callus. Among these, 7.4 and 6.6 million were unique reads aligned to the reference sequences (Table 1). Unique sRNAs ranged from 18 to 30 nt in length in the two libraries (Fig. 3). The most common lengths of unique sequences in each library were 21–24 nt, with 24 nt long reads being the majority, followed by 23 nt.

**Fig. 3.**
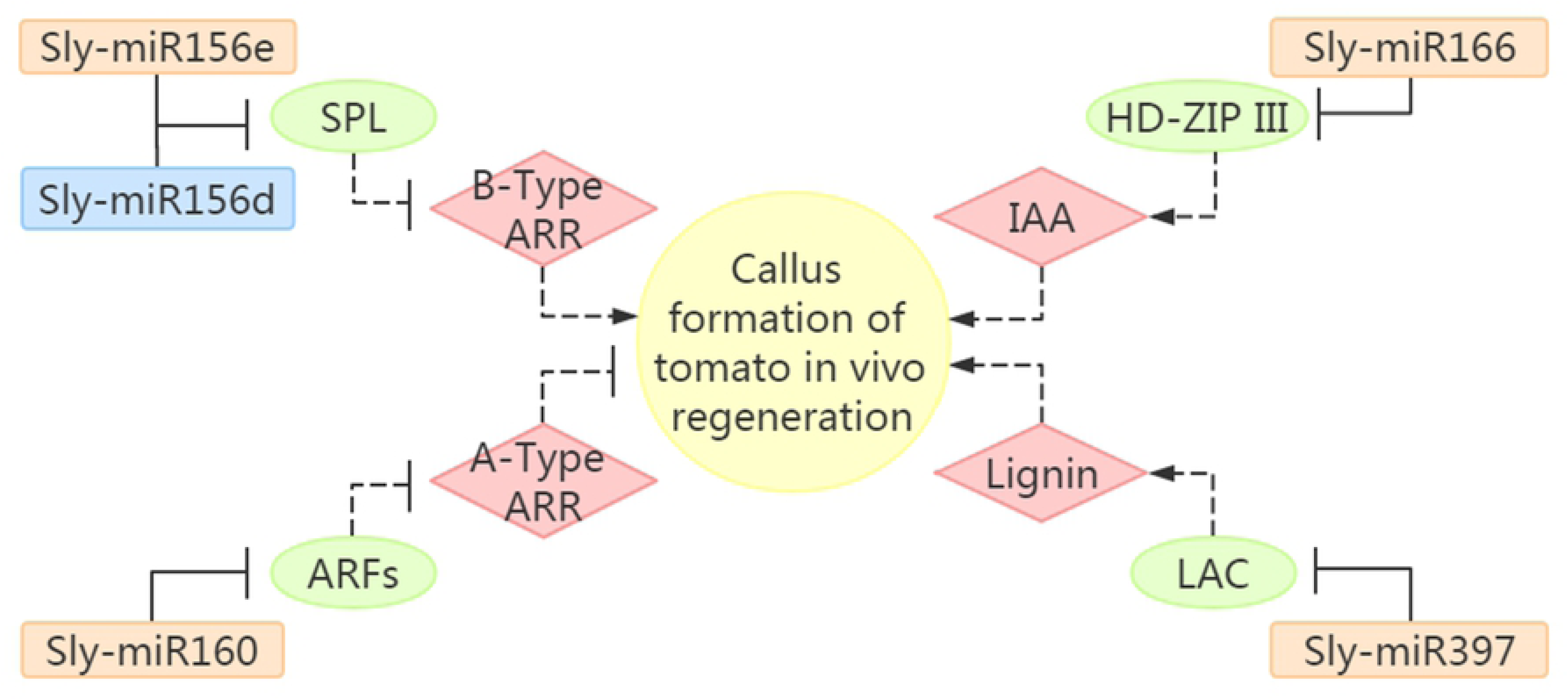
Length distributions of unique sRNAs in stem and callus.

**Table 1.**
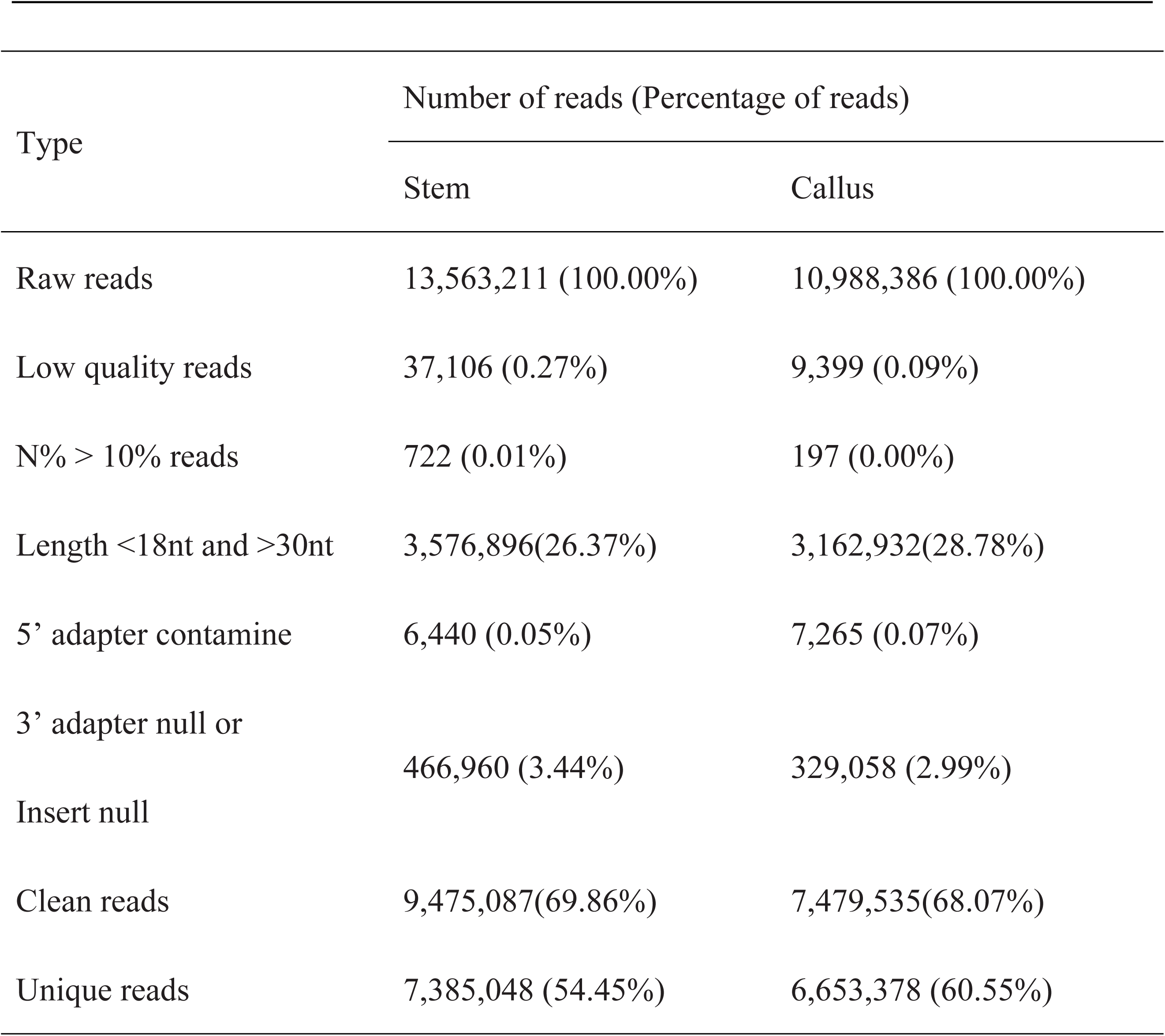
Sequencing data filtering of two sRNA libraries produced from stem and callus.

Table 2 summarizes the categories of unique reads. High levels of small RNA expression from rRNA and NAT genes were observed in both libraries. The number of miRNAs was more abundant in stem tissue as compared to the callus of tomato, mainly due to the high expression of sly-miR171, sly-miR396 and sly-miR397.

**Table 2.**
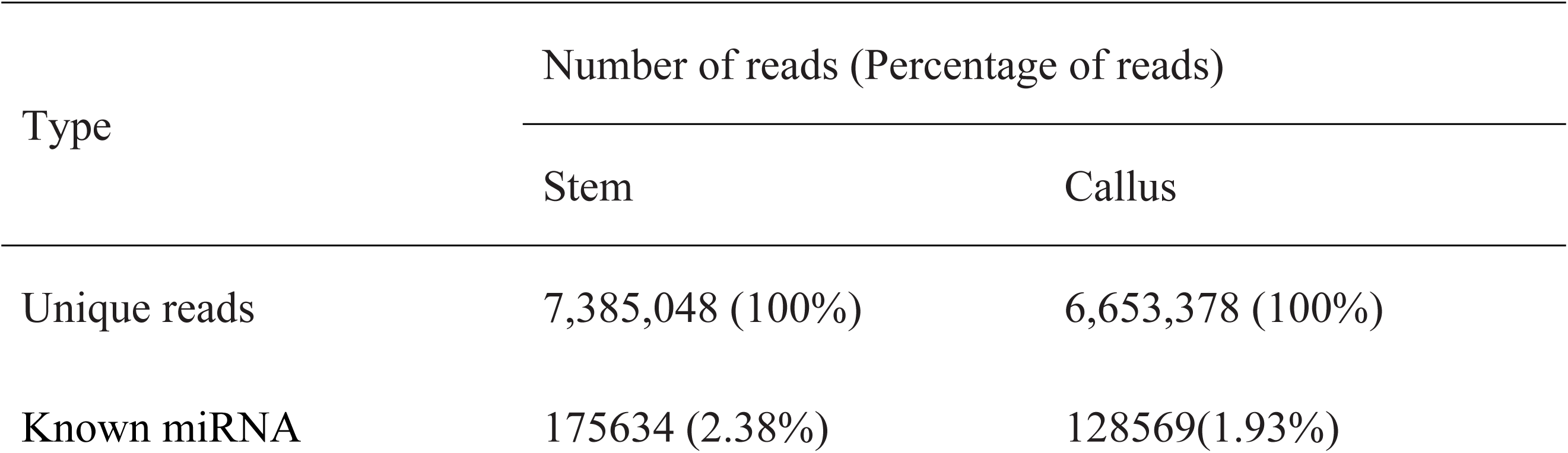
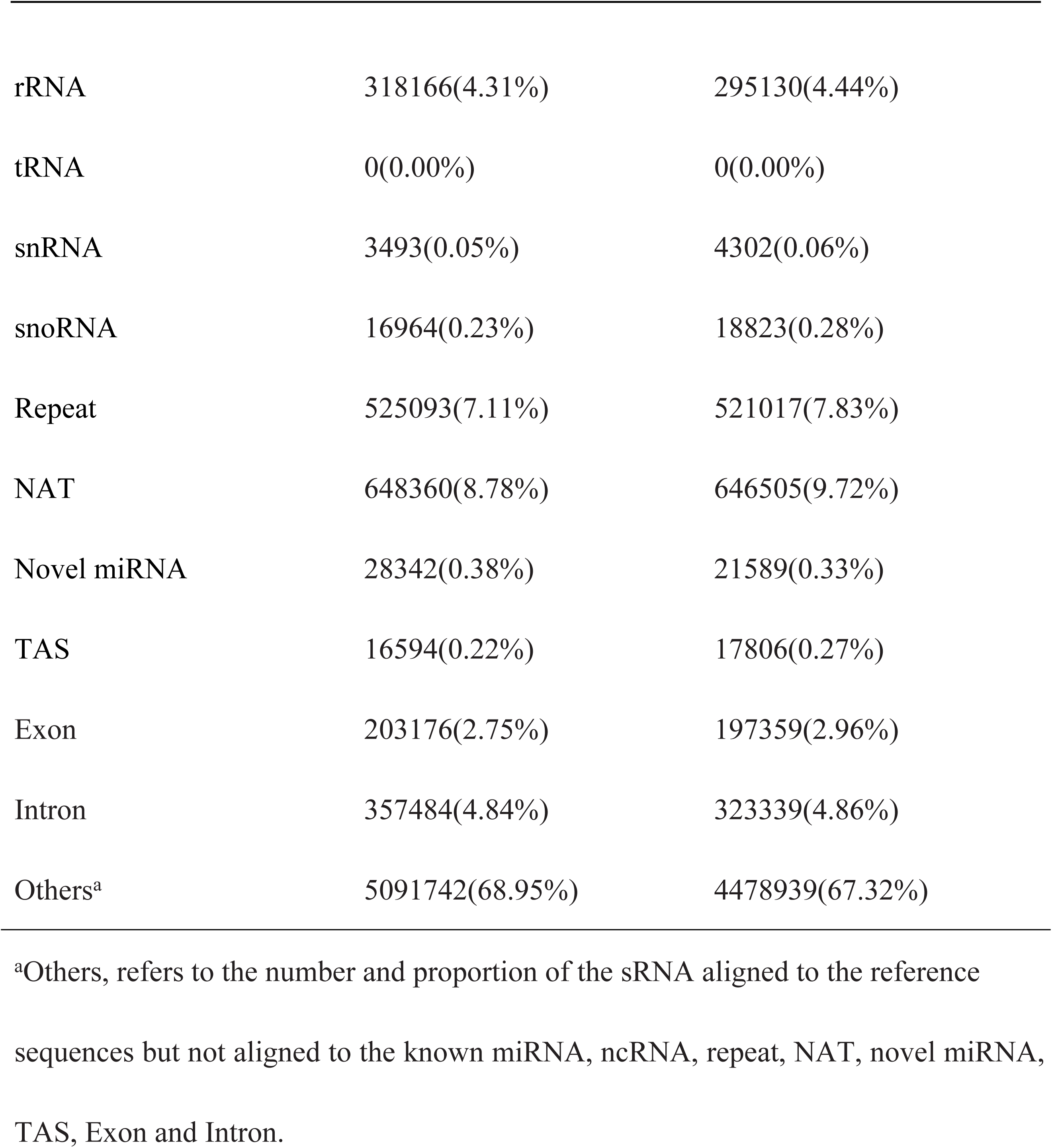
Reads categories of two small RNA libraries derived from stem and callus.

| Type | Number of reads (Percentage of reads) |  |
| --- | --- | --- |
|  | Stem | Callus |
| Unique reads | 7,385,048 (100%) | 6,653,378 (100%) |
| Known miRNA | 175634 (2.38%) | 128569(1.93%) |
| rRNA | 318166(4.31%) | 295130(4.44%) |
| tRNA | 0(0.00%) | 0(0.00%) |
| snRNA | 3493(0.05%) | 4302(0.06%) |
| snoRNA | 16964(0.23%) | 18823(0.28%) |
| Repeat | 525093(7.11%) | 521017(7.83%) |
| NAT | 648360(8.78%) | 646505(9.72%) |
| Novel miRNA | 28342(0.38%) | 21589(0.33%) |
| TAS | 16594(0.22%) | 17806(0.27%) |
| Exon | 203176(2.75%) | 197359(2.96%) |
| Intron | 357484(4.84%) | 323339(4.86%) |
| Others<sup>a</sup> | 5091742(68.95%) | 4478939(67.32%) |

<sup>a</sup>Others, refers to the number and proportion of the sRNA aligned to the reference sequences but not aligned to the known miRNA, ncRNA, repeat, NAT, novel miRNA, TAS, Exon and Intron.

### Identification of known miRNAs

To identify known miRNAs, sRNA sequences obtained from deep sequencing were contrasted to other currently annotated miRNAs of known mature plant species in miRBase. A total of 92 known miRNAs were identified, belonging to 29 miRNA gene families in the two sRNA libraries. Overall, 88 and 91 mature miRNAs were identified in the stem and callus tissues, respectively (S3 Table). As shown in Fig. 4, the sly-miR159 family was the most abundantly expressed, while sly-miR9471, sly-miR6022, sly-miR396, and sly-miR166 families were moderately abundant. Furthermore, the secondary structures of known miRNAs are shown in Fig. 5 (A) and S1 Fig.

**Fig. 4.**
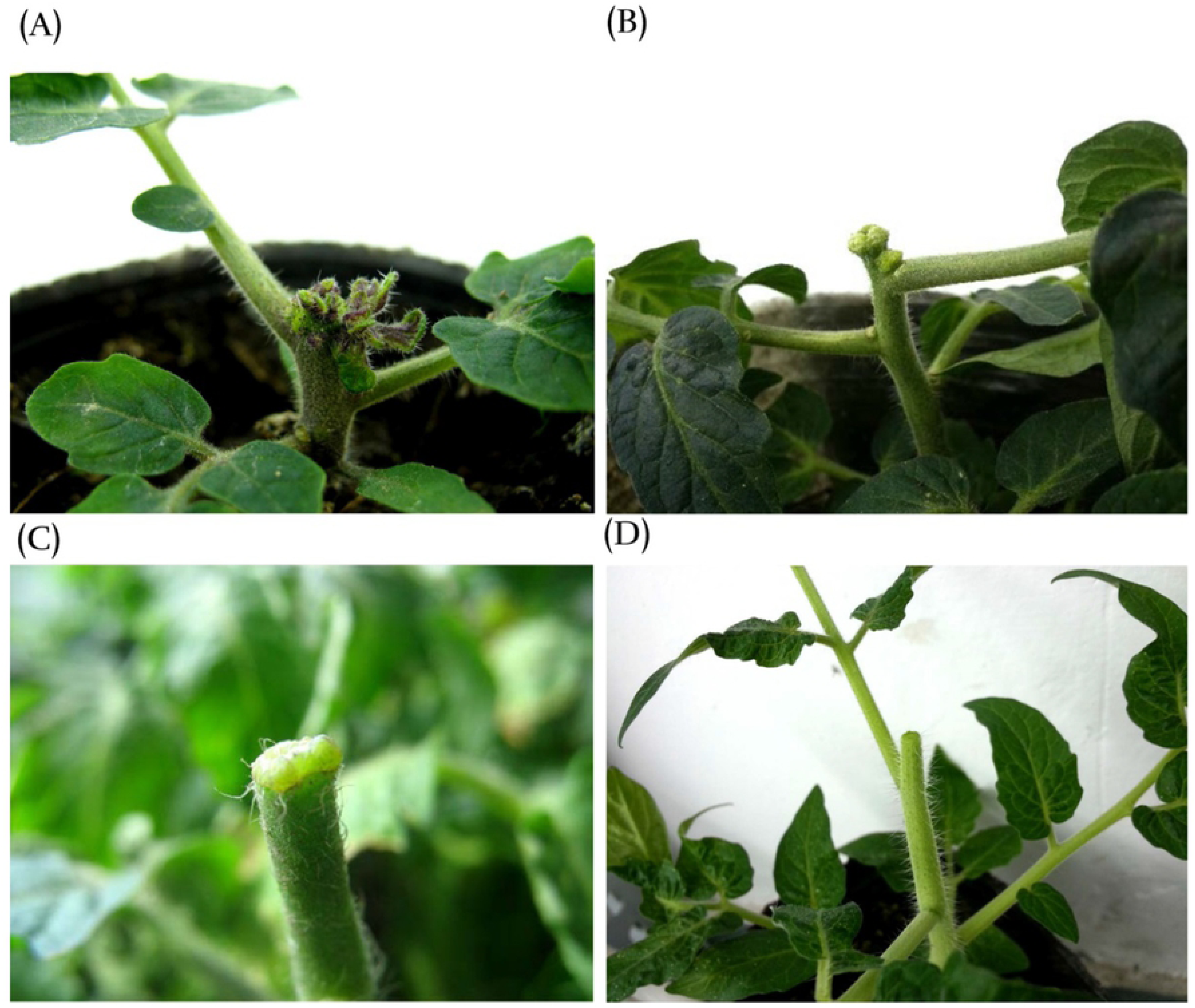
Reads of known miRNA families at stem and callus.

**Fig. 5.**
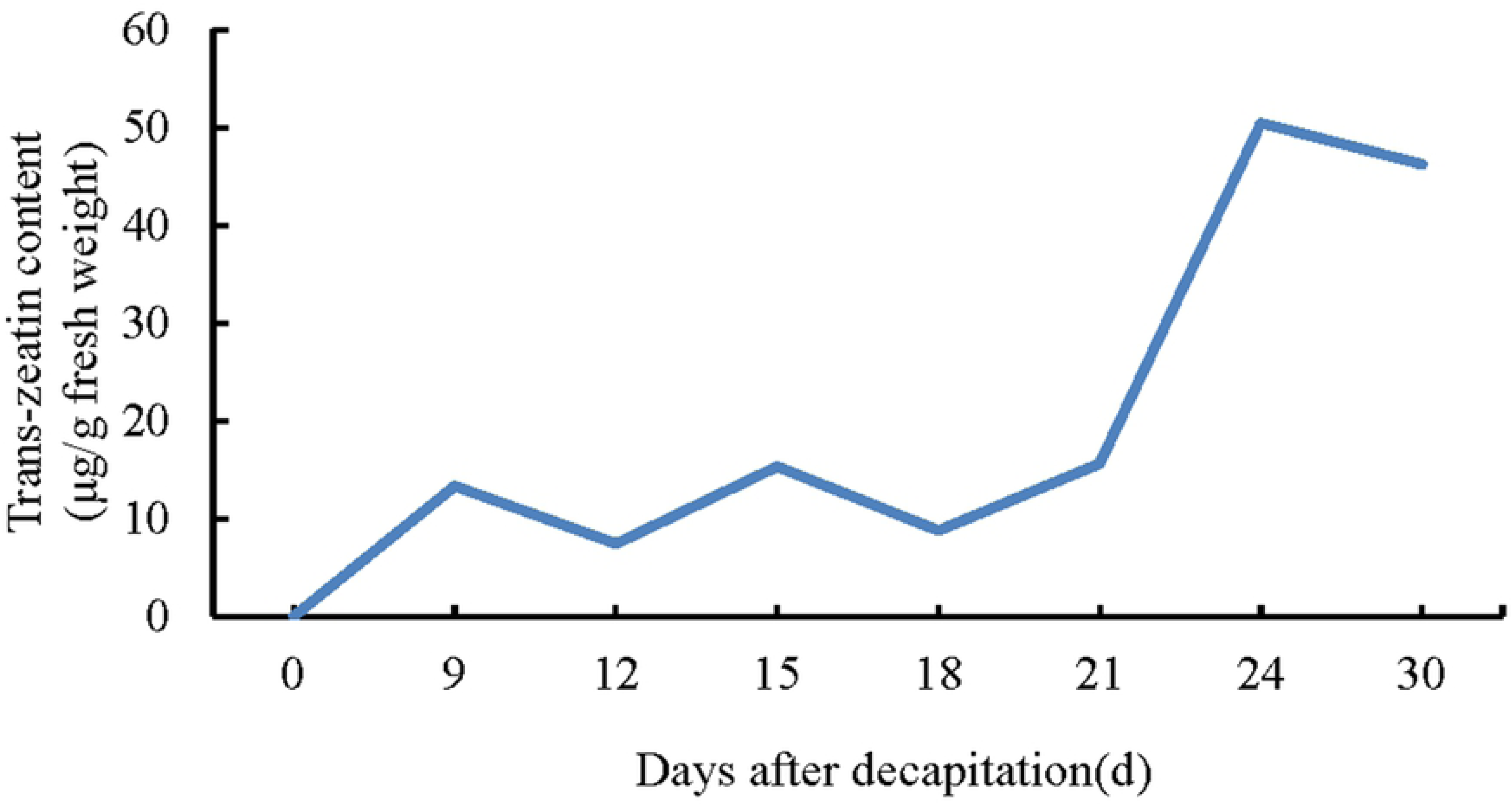
Secondary structure of identified miRNA precursors. The red protrusions are the mature sequences. (A) Known miRNA: sly-miR159; (B) Novel miRNA: novel 110.

### Predicted novel miRNAs

Unannotated miRNAs were used to predict novel miRNAs. We identified 91 novel miRNAs were identified in total, of which 82 were mapped in both libraries (S4 Table). The expression levels of novel miRNAs were distinctly different. Most of them showed comparatively low expression levels (63 novel miRNAs in stem samples and 68 novel miRNAs in callus had less than 120 raw reads). In contrast, two novel miRNAs (annotated as novel 1 and novel 9) in both libraries contained more than 1,000 reads. The most abundantly expressed novel miRNA was novel 1 with a total of 23,938 reads in both libraries. The predicted secondary structures of novel pre-miRNAs are showed in Fig. 5 (B) and S2 Fig.

### Identification of differentially expressed miRNAs

A total of 49 known and 44 novel miRNAs pertaining to the two libraries were expressed with significant differences with regards to log_2_ (callus/stem) (>1 or<-1) and P-value (<0.05) criteria (Fig. 6) (S5 Table). For known miRNAs, 24 miRNAs were up-regulated and 25 miRNAs were down-regulated in callus vs. stem tissue samples. Among the novel miRNAs, 17 were up-regulated and 27 were down-regulated in callus vs. stem tissues (Fig. 7). When miRNA distributions were assessed between the two libraries, 44 known defined miRNAs and 37 novel miRNAs were generally expressed in both libraries. The comparison of miRNA expression showed that 1 known and 7 novel miRNAs were expressed only in the stem, while 4 novel miRNAs were expressed solely in callus tissue, respectively (Fig. 8).

**Fig. 6.**
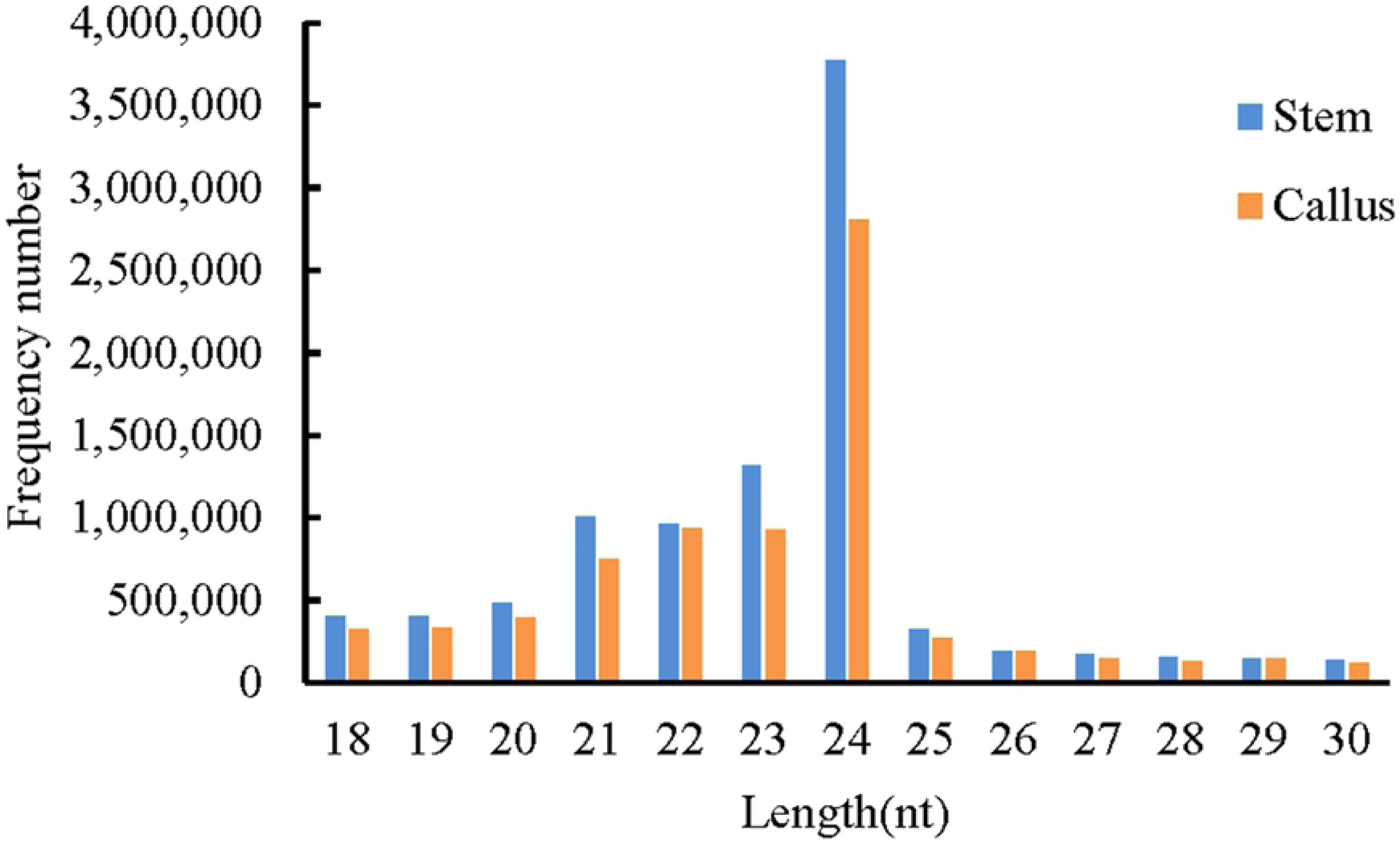
Cluster analyses of differentially expressed miRNAs. Red denotes highly expressed miRNAs, while blue denotes weakly expressed miRNAs. The color is from red to blue, indicating that log_10_ (TPM + 1) is from large to small.

**Fig. 7.**
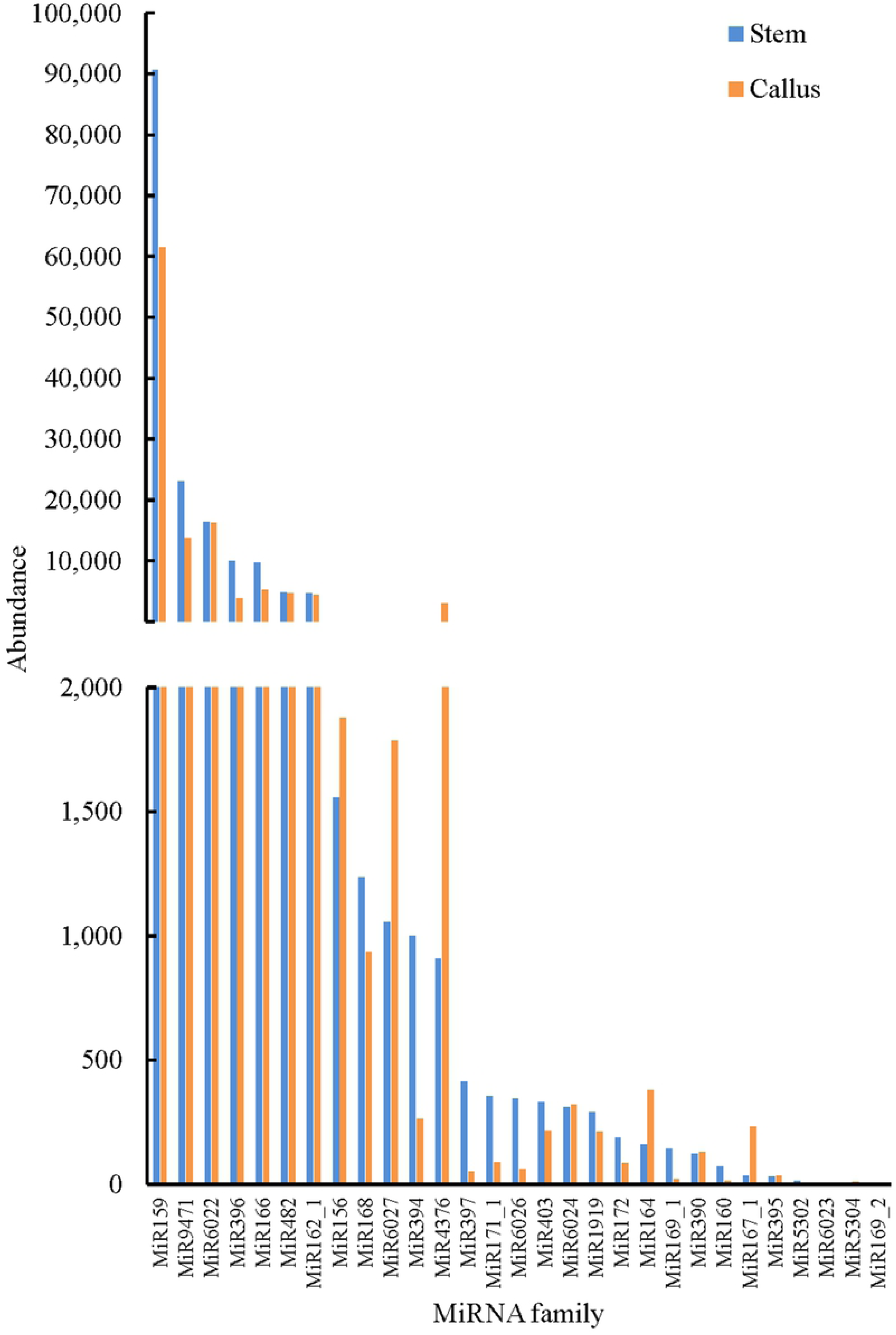
The number of known and novel up- and down-regulated miRNAs in callus vs. stem tissue.

**Fig. 8.**
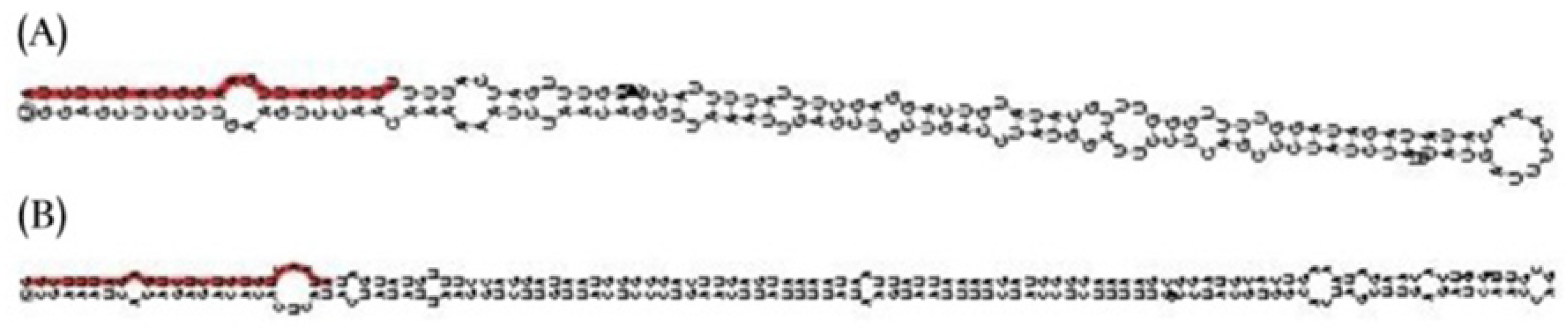
Venn diagram of the number of specifically expressed miRNAs at stem and callus. (A) Known miRNAs; (B) Novel miRNAs.

The miRNAs with lower p-value include sly-miR166 and sly-miR397. The significantly down-regulated expression of sly-miR166 in callus cells could be related to its role in promoting callus formation by down-regulating homeodomain leucine zipper class III (HD-ZIP III) levels [49–51]. The most significantly down-regulated gene in callus tissue was sly-miR397, which is known to play an important role in the accumulation of laccases during callus formation[52, 53].

To confirm miRNAs expression levels in stem and callus and verify the deep-sequencing results, four known and two novel miRNAs were selected randomly for qRT-PCR. These miRNAs expression patterns resembled the deep-sequencing results, suggesting that sRNA sequencing data were reliable (Fig. 9).

**Fig. 9.**
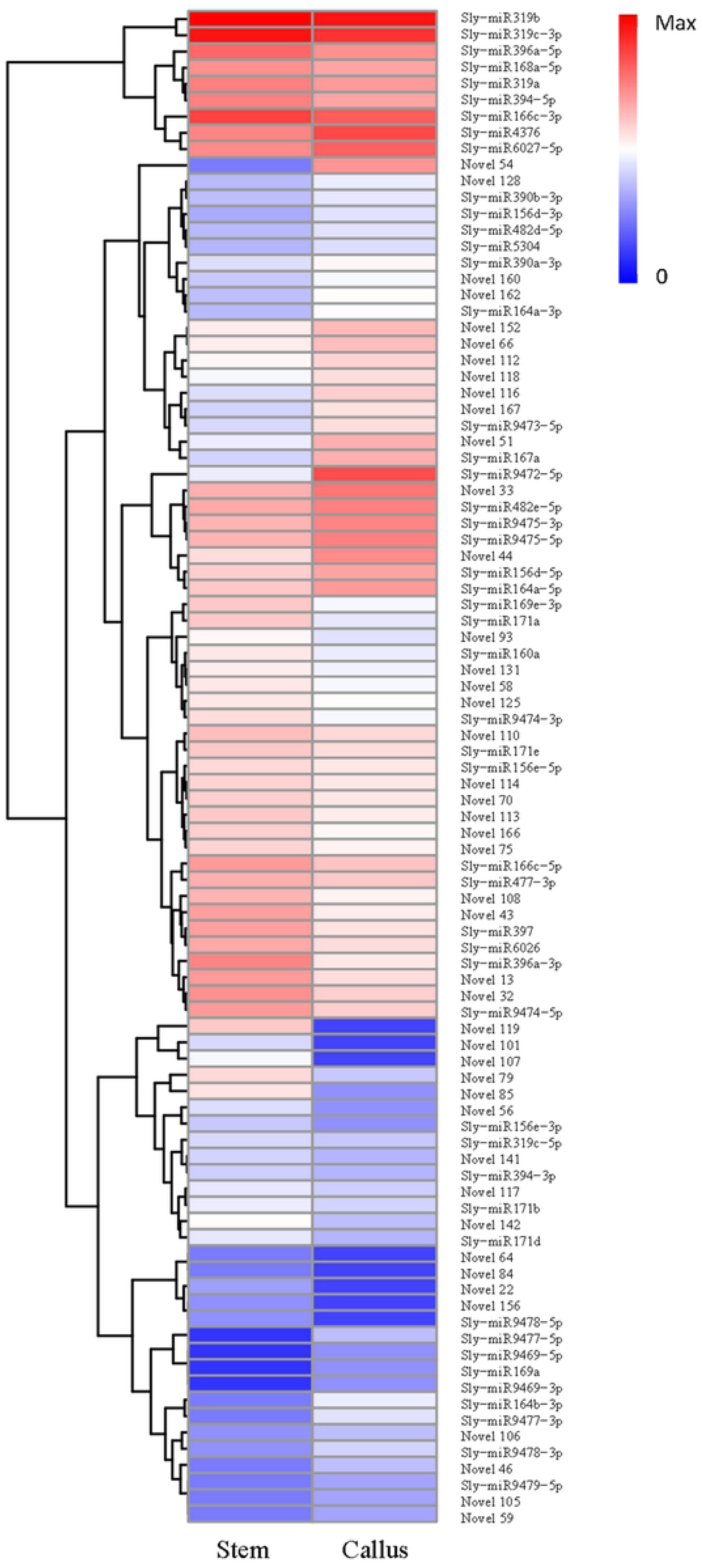
The relative expression levels of 6 (four known and two novel) miRNAs by qRT-PCR. The bars represents the relative expression and standard deviation of the 6 miRNAs. qRT-PCR value of miRNAs in stem was set to 1, and values of miRNAs in callus were scaled.

### Target prediction

To analyze the biological functions of differentially expressed miRNAs in stem and callus tissues, the psRobot software was used to predict target genes. Among 1186 predicted target genes, a total of 505 known and 6 novel differentially expressed miRNA target genes were identified (S6 Table). Functional annotations of BLAST analysis for predicted target genes indicated that these targets contained mRNA coding regions for zinc finger protein (sly-miR165, sly-miR164, sly-miR391, sly-miR394, sly-miR396, sly-miR477, sly-miR482, sly-miR6027, sly-miR9469, sly-miR9478, and sly-miR9479), and MYB (sly-miR156, sly-miR319, sly-miR9469, and sly-miR9478) protein. Furthermore, some miRNAs were found to target transcription factors, such as SQUAMOSA promoter binding protein-like gene (SPL) (sly-miR156)[54], Auxin response factors (ARFs) (sly-miR160)[55], HD-ZIP III (sly-miR166)[56], NAM (sly-miR164 and sly-miR9478)[57], and MADS (sly-miR396 and sly-miR9477)[58], which are all known to be involved in plant regeneration. Laccase(sly-miR397) was also important in the regulation of plant development and regeneration[22, 23]. The target genes of some miRNAs specifically expressed in the callus were CCAAT-binding (sly-miR169a), zinc finger (sly-miR9469-3p and sly-miR9469-5p), SQUAMOSA promoter binding protein (SBP-box), MADS-box and K-box (sly-miR9477-5p). Interestingly, all the target genes of novel 46 (solyc05g015840.2, solyc12g038520.1, solyc10g078700.1, solyc05g015510.2, solyc05g012040.2, and solyc04g045560.2) were the same as those targeted by sly-miR156e-5p, sly-miR156d-5p, and sly-miR156a.

## Discussion

Recently great progress has been made in understanding the role of miRNAs in regulating the transitions between different development stage in plants, such as those from vegetative-to-reproductive, juvenile-to-adult and aerial stem-to-rhizome transitions[59–62]. In the present study, we demonstrate that miRNAs are involved in complex regulatory networks during stem-callus transition during *in vivo* regeneration of tomato. We identified a total of 183 miRNAs (92 conserved and 91 novel miRNAs) by next-generation sequencing. Previous studies on miRNAs, including Xu et al.[30] identified 289 known miRNAs and 1087 novel miRNAs in longan, while Wu et al. [26] reported 50 known and 45 novel miRNAs in citrus. Taken together, these data show that distinct types of miRNAs are expressed at different levels during the process of regeneration in different species.

Cytokinin triggers a complex gene expression program in plant tissue culture that results in adventitious shoot regeneration[63]. The current model for cytokinin signal transduction is a multi-step phosphorelay. First, Arabidopsis histidine kinase (AHKs), the cytokinin receptors in the plasma membrane, perceive the cytokinin signal triggering a multi-step phosphorelay. At the end of this pathway, B-ARR receives the phosphoryl group and becomes active. As transcription factors, B-type ARRs can activate the expression of cytokinin-responsive genes and A-type ARRs. Interestingly, the expression of A-type ARRs interferes with the function of B-type ARR proteins through a negative feedback loop[64, 65]. Cytokinin thus plays a vital role during *in vitro* regeneration. It can not only induce adventitious buds alone, but also cooperate with auxin. Many studies have confirmed that miRNA regulate hormone signaling genes involved in regeneration.

### MiR156/SPL module involved in callus formation by regulating cytokinin signaling pathway

Siddiqui et al. [66] summarized the most common expressed miRNAs during SEG in 11 economically plants. Of these, miRNA156 was found to be most frequently detected in six of the 11 plant species tested. Sequences of miR156 were highly conserved in plants [67]. In the present study, sly-miR156d-5p and sly-miR156e-5p were newly identified and shown to be expressed differently in stem and callus tissue. Promoter binding protein (SBP) domain SQUAMOSA was predicted to be one of the targets of sly-miR156d-5p and sly-miR156e-5p. SBP domain proteins, putative plant-specific transcription factor gene families, have been shown to participate in various plant biological processes and to be involved in vegetative-to-reproductive phase transition[68–70], pollen sac development[71], gibberellins (GAs) signaling network [72] and establishment of lateral meristems[73]. As SBP-box gene family members, 10 of 16 *SPL* genes were shown to be targets of miR156 in *Arabidopsis*, while 10 of the 15 *SPL* genes were proposed to be targets of miR156 in citrus [74, 75]. The function of the miR156-SPLs module was confirmed to be crucial in callus production in citrus *in vitro* callus through targeted inhibition of miR156-targeted SPLs and over-expression of csi-miR156a[20]. Therefore, differential expression levels of miR156 during tomato callus generation in the present study, suggest that is likely to play an important role in *in vivo* regeneration.

Zhang et al. [76] demonstrated that miR156 participates in regulation of shoot regeneration *in vitro*. MiR156 expression gradually increases with age and suppresses the expression of its target *SPL* genes. Down-regulated SPLs attenuate cytokinin signaling by binding to the B-type Arabidopsis response regulators (ARR) transcription factor. The data presented here show that cytokinin levels increase during *in vivo* regeneration in tomato. However, sly-miR156d-5p and sly-miR156e-5p were found to be up- and down-regulated, respectively. Thus, the regulation of miR156-SPL-ARR module during *in vivo* callus formation and shoot regeneration in tomato needs to be further investigated.

### IAA level regulated by miR166 in callus formation

Low expression levels of sly-miR166c-5p and sly-miR166c-3p were observed during the change from stem to callus stages in this study. Previous research has shown that miR166, together with miR156 and miR396 were down-regulated during callus formation from tea plant stem explants[77]. miR166 was identified to target Class III homeodomain leucine zipper (HD-Zip III) gene family of transcription factors, including REVOLUTA (REV), PHABULOSA (PHB), PHAVOLUTA (PHV), CORONA (CNA), and ATHB8 in *Arabidopsis* [49]. HD-ZIP III proteins play an important role in plant regeneration by regulating the differentiation of stem cells and the establishment of shoot apical meristem (SAM) and RAM [78–80].

More recently, REV was demonstrated to activate genes upstream of several auxin biosynthesis, transport, and response genes. Brandt et al. [81] identified that REV targeted the auxin biosynthetic enzymes TAA1 and YUCCA5(YUC5), and directly affected the levels of free auxin. In *Arabidopsis*, loss-of-function mutants of REV showed lower expression levels of the PIN1 and PIN2 auxin transporters and reduction in the tip-to-base transport of auxin[82]. Additionally, REV function is necessary for polar auxin transport in the shoot[83]. Li et al.[51,84,85] demonstrated that over-expression of LaMIR166a and down-regulated of LaHDZIP31-34 genes results in different IAA levels in pro-embryogenic masses of *L. leptolepis*. The authors speculated that LaMIR166 targeted HD-ZIP III genes likely regulate auxin biosynthesis and response genes. Overall, these results indicated the complex regulaory relationships between miR166 and plant development. Further, Ma et al. [62] reviewed five key microRNAs involved in developmental phase transitions in seed plant, and miR166 was one of them.

### Other miRNAs related to the phytohormone signaling during in vivo regeneration

There is no doubt that auxin signaling and transport is a versatile trigger of plant developmental changes incluing regeneration [86]. Based on a number of previous studies, which focussed on miRNAs involved in regulation of the auxin signaling, in *Arabidopsis*, miR393 was shown to contribute to SE, leaf development and antibacterial resistance by repressing auxin signaling[87–89]. Two auxin response factors genes, *ARF6* and *ARF8*, are targeted by miR167 [90]. During callus formation, miR160 was defined as a key repressor by modulating the interplay between auxin and cytokinin. The callus initiation was repressed by over-expression of miR160 or reduced expression of its target *ARF10*. *ARF10* can inhibit cytokinin signaling A-type genes *ARR15*[91]. Although A-type genes *ARR15* and *ARR7* were identified to inhibit callus formation, those of the B-type genes *ARR1* and *ARR21* can enhance its initiation[92, 93](Fig. 10).

**Fig. 10.**
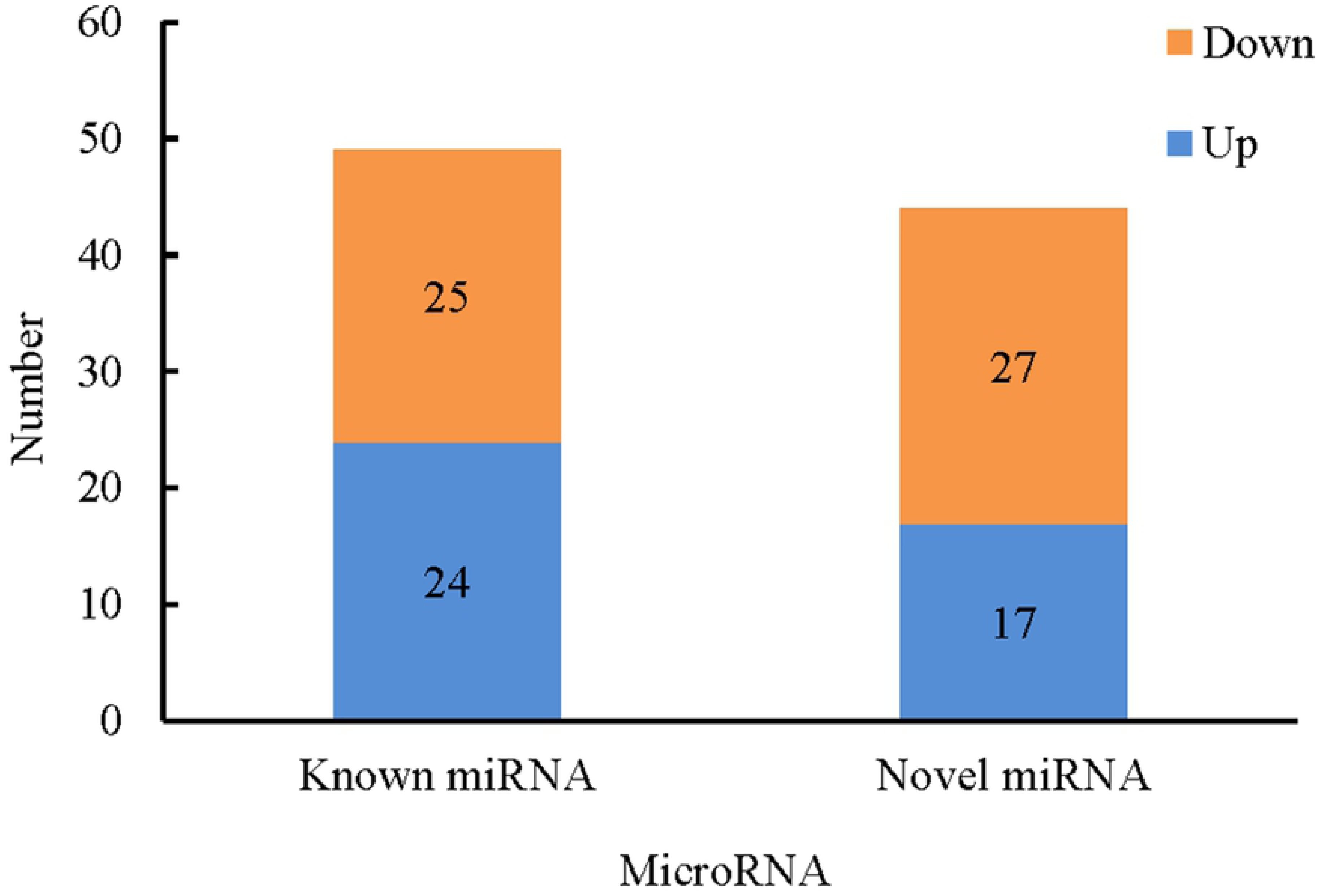
Genetic networks of callus formation during *in vivo* regeneration of tomato regulated by miRNA-target modules together with their downstream targets. Arrows represent activation, while lines with a bar represent repression. The solid lines represent the results predicted by this study, and the dotted lines represent the results from references. The up-regulated miRNAs are shown in the blue box, while the down-regulated miRNAs are shown in orange ones. miRNA targets are shown in green oval frames.

Cell proliferation relied not only on high levels of auxin but also on low level of cytokinin during in vitro callus induction in *Arabidopsis* [94]. This study showed the down-regulation of sly-miR160 and low concentrations of cytokinin in callus were crucial in callus formation. We predict that a similar interplay between microRNA/phytohormone levels may exist between *in vivo* and *in vitro* regeneration in tomato.

### MiR397 repressed callus formation through inhibition of laccases expression

The expression of Sly-miR397 was most significantly down-regulated in callus tissue. MiR397 has been validated to target laccases (*LAC2* and *LAC17* in this study), a group of polyphenol oxidases[52, 95]. In higher plants, laccases are associated with lignin and xylem synthesis, and are proposed to play a role in secondary cell wall thickening [23,96,97]. Lignin is an essential component of plant secondary cell walls, which influence plant growth and differentiation[98]. Previous studies indicated that callus tissue first contains lignified parenchyma cells, followed by the formation of short vessels and traumatic resin ducts after plant injury, and the induction of vessels requires the involvement of lignin[53, 99]. Overall, we predict that low expression levels of *Sly-miR397* in callus tissue permits the accumulation of laccases, leading to the increase of lignin deposition within the callus.

## Conclusion

We used a new model system to study the dynamic changes in trans-zeatin levels and the regulatory patterns of miRNA expression during *in vivo* regeneration of tomato. The significant changes in trans-zeatin levels at 0, 9, 12, 15, 18, 21, 24 and 30 d after decapitation proves that trans-zeatin plays a crucial role during *in vivo* regeneration in tomato. However, the treatment with excess lovastatin on the cut surface of tomato stems did not inhibit callus formation, which indicated that *de novo* biosynthesis of cytokinin did not occur in the cut surface of tomato stems. A total of 92 known and 91 novel miRNAs were identified from the stem explant and the callus regenerated from the cutting surfaces after decapitation, respectively, of which 49 known and 44 novel miRNAs exhibited differential expression between the two libraries. In addition, a total of 505 known miRNA target genes and 6 novel miRNA target genes were further identified. We predict that these differentially expressed miRNAs and their relevant target genes play an important role in callus formation during *in vivo* regeneration of tomato. Among these, sly-miR156, sly-miR160, sly-miR166, and sly-miR397 are predicted to be involved in callus formation during *in vivo* regeneration of tomato by targeting SPL, HD-ZIP III, ARFs, and LAC proteins, as well as by regulating cytokinin, IAA, and laccase levels. The findings of this study provide a useful resource for further investigation on callus formation during *in vivo* regeneration of tomato.

## Supporting information captions

**S1 Fig. The secondary structures of known miRNAs.** The whole sequences are miRNA precursors, and the red prominent parts are the mature sequences.

**S2 Fig. The secondary structures of novel pre-miRNAs.** The whole sequences are miRNA precursors, and the red prominent parts are the mature sequences.

**S1 Table. Primers used in this study for qRT-PCR.**

**S2 Table. Quality of raw reads of two sRNA libraries produced from stem and callus.**

^a^ Q20, The percentage of bases with Phred value greater than 20 in the total bases.

^b^ Q30, The percentage of bases with Phred value greater than 30 in the total bases

**S3 Table. Detailed information of known miRNAs in two small RNA libraries derived from stem and callus.**

**S4 Table. Novel miRNAs of two small RNA libraries derived from stem and callus.**

**S5 Table. Differentially expressed known and novel miRNAs during callus formation of tomato *in vivo* regeneration.**

**S6 Table. Target genes for differentially expressed known and novel miRNAs.**

## References

1. Altpeter F, Springer NM, Bartley LE, Blechl A, Brutnell TP, Citovsky V, et al. Advancing Crop Transformation in the Era of Genome Editing. Plant Cell. 2016; tpc.00196.2016. doi:10.1105/tpc.16.00196

2. Maher MF, Nasti RA, Vollbrecht M, Starker CG, Clark MD, Voytas DF. Plant gene editing through de novo induction of meristems. Nat Biotechnol. 2020;38: 84– 89. doi:10.1038/s41587-019-0337-2

3. Zuo J, Niu Q-W, Frugis G, Chua N-H. The WUSCHEL gene promotes vegetative-to-embryonic transition in Arabidopsis. Plant J. 2002;30: 349–359. doi:10.1046/j.1365-313x.2002.01289.x

4. Barton MK. Twenty years on: The inner workings of the shoot apical meristem, a developmental dynamo. Developmental Biology. 2010;341: 95–113. doi:10.1016/j.ydbio.2009.11.029

5. Lowe K, Wu E, Wang N, Hoerster G, Hastings C, Cho M-J, et al. Morphogenic Regulators Baby boom and Wuschel Improve Monocot Transformation. Plant Cell. 2016;28: 1998–2015. doi:10.1105/tpc.16.00124

6. Nelson-Vasilchik K, Hague J, Mookkan M, Zhang ZJ, Kausch A. Transformation of Recalcitrant Sorghum Varieties Facilitated by Baby Boom and Wuschel2. Curr Protoc Plant Biol. 2018;3: e20076. doi:10.1002/cppb.20076

7. Yin L, Lan Y, Zhu L. Analysis of the protein expression profiling during rice callus differentiation under different plant hormone conditions. Plant Mol Biol. 2008;68: 597–617. doi:10.1007/s11103-008-9395-4

8. Rodrigues AS, Chaves I, Costa BV, Lin Y-C, Lopes S, Milhinhos A, et al. Small RNA profiling in Pinus pinaster reveals the transcriptome of developing seeds and highlights differences between zygotic and somatic embryos. Sci Rep. 2019;9: 11327. doi:10.1038/s41598-019-47789-y

9. Amutha S, Kathiravan K, Singer S, Jashi L, Shomer I, Gaba SV. Adventitious Shoot Formation in Decapitated Dicotyledonous Seedlings Starts with Regeneration of Abnormal Leaves from Cells Not Located in a Shoot Apical Meristem. Vitro Cellular & Developmental Biology Plant. 2009;45: 758–768. doi:10.2307/25623037

10. Masafumi J, Genjirou Mori, Kazuhiko Mitsukuri. In Vivo Shoot Regeneration Promoted by Shading the Cut Surface of the Stem in Tomato Plants. Hortscience A Publication of the American Society for Horticultural Science. 2008;43: 220–222. doi:10.21273/HORTSCI.43.1.220

11. Nielsen MD, Farestveit B, Andersen AS. Adventitious Shoot Development from Decapitated Plants of Periclinal Chimeric Poinsettia Plants (Euphorbia pulcherrima Willd ex Klotsch). European Journal of Horticultural Science. 2003 [cited 12 Apr 2020]. Available: http://www.jstor.org/stable/24126204

12. Jones-Rhoades MW, Bartel DP, Bartel B. MicroRNAs AND THEIR REGULATORY ROLES IN PLANTS. Annual Review of Plant Biology. 2006;57: 19–53. doi:10.1146/annurev.arplant.57.032905.105218

13. Man W, Fei Y, Wei W, Yu W, Han D, Qian L. Epstein-Barr virus-encoded microRNAs as regulators in host immune responses. International Journal of Biological Sciences. 2018;14: 565–576. doi:10.7150/ijbs.24562

14. Lima JCD, Loss-Morais G, Margis R. MicroRNAs play critical roles during plant development and in response to abiotic stresses. Genetics & Molecular Biology. 2012;35: 1069–1077. doi:10.1590/S1415-47572012000600023

15. Li C, Zhang B. MicroRNAs in Control of Plant Development. J Cell Physiol. 2016;231: 303–313. doi:10.1002/jcp.25125

16. Smoczynska A, Szweykowska-Kulinska Z. MicroRNA-mediated regulation of flower development in grasses. Acta Biochim Pol. 2016;63: 687–692. doi:10.18388/abp.2016_1358

17. Singh A, Gautam V, Singh S, Sarkar Das S, Verma S, Mishra V, et al. Plant small RNAs: advancement in the understanding of biogenesis and role in plant development. Planta. 2018;248: 545–558. doi:10.1007/s00425-018-2927-5

18. Chávez-Hernández EC, Alejandri-Ramírez ND, Juárez-González VT, Dinkova TD. Maize miRNA and target regulation in response to hormone depletion and light exposure during somatic embryogenesis. Front Plant Sci. 2015;6: 555. doi:10.3389/fpls.2015.00555

19. Su YH, Liu YB, Zhou C, Li XM, Zhang XS. The microRNA167 controls somatic embryogenesis in Arabidopsis through regulating its target genes ARF6 and ARF8. Plant Cell Tiss Organ Cult. 2016;124: 405–417. doi:10.1007/s11240-015-0903-3

20. Long JM, Liu CY, Feng MQ, Yun L, Wu XM, Guo WW. miR156-SPL modules regulate induction of somatic embryogenesis in citrus callus. Journal of Experimental Botany. 2018; 12. doi:10.1093/jxb/ery132

21. Chu Z, Chen J, Xu H, Dong Z, Chen F, Cui D. Identification and Comparative Analysis of microRNA in Wheat (Triticum aestivum L.) Callus Derived from Mature and Immature Embryos during In vitro Culture. Frontiers in Plant Science. 2016;7. doi:10.3389/fpls.2016.01302

22. Luo Y-C, Zhou H, Li Y, Chen J-Y, Yang J-H, Chen Y-Q, et al. Rice embryogenic calli express a unique set of microRNAs, suggesting regulatory roles of microRNAs in plant post-embryogenic development. FEBS Letters. 2006;580: 5111–5116. doi:10.1016/j.febslet.2006.08.046

23. Chen CJ, liu Q, Zhang Y-C, Qu L-H, Chen Y-Q, Gautheret D. Genome-wide discovery and analysis of microRNAs and other small RNAs from rice embryogenic callus. Rna Biology. 2011;8: 538–547. doi:10.4161/rna.8.3.15199

24. Yang X, Wang L, Yuan D, Lindsey K, Zhang X. Small RNA and degradome sequencing reveal complex miRNA regulation during cotton somatic embryogenesis. Journal of Experimental Botany. 2013;64: 1521–1536. doi:10.1093/jxb/ert013

25. Gao C, Wang P, Zhao S, Zhao C, Xia H, Hou L, et al. Small RNA profiling and degradome analysis reveal regulation of microRNA in peanut embryogenesis and early pod development. Bmc Genomics. 2017;18: 220. doi:10.1186/s12864-017-3587-8

26. Wu XM, Kou SJ, Liu YL, Fang YN, Xu Q, Guo WW. Genomewide analysis of small RNAs in nonembryogenic and embryogenic tissues of citrus: microRNA- and siRNA-mediated transcript cleavage involved in somatic embryogenesis. Plant Biotechnology Journal. 2015;13: 383–394. doi:10.1111/pbi.12317

27. Sabana AA, Rajesh MK, Antony G. Dynamic changes in the expression pattern of miRNAs and associated target genes during coconut somatic embryogenesis. Planta. 2020;251: 79. doi:10.1007/s00425-020-03368-4

28. Zhang J, Zhang S, Han S, Wu T, Li X, Li W, et al. Genome-wide identification of microRNAs in larch and stage-specific modulation of 11 conserved microRNAs and their targets during somatic embryogenesis. Planta. 2012;236: 647–657. doi:10.1007/s00425-012-1643-9

29. Yakovlev IA, Fossdal CG. In Silico Analysis of Small RNAs Suggest Roles for Novel and Conserved miRNAs in the Formation of Epigenetic Memory in Somatic Embryos of Norway Spruce. Front Physiol. 2017;8: 674. doi:10.3389/fphys.2017.00674

30. Xu X, Chen X, Chen Y, Zhang Q, Su L, Chen X, et al. Genome-wide identification of miRNAs and their targets during early somatic embryogenesis in Dimocarpus longan Lour. Sci Rep. 2020;10: 4626. doi:10.1038/s41598-020-60946-y

31. Li T, Chen J, Qiu S, Zhang Y, Wang P, Yang L, et al. Deep sequencing and microarray hybridization identify conserved and species-specific microRNAs during somatic embryogenesis in hybrid yellow poplar. PLoS ONE. 2012;7: e43451. doi:10.1371/journal.pone.0043451

32. Zhai L, Xu L, Wang Y, Huang D, Yu R, Limera C, et al. Genome-Wide Identification of Embryogenesis-Associated microRNAs in Radish (Raphanus sativusL.) by High-Throughput Sequencing. Plant Molecular Biology Reporter. 2014;32: 900–915. doi:10.1007/s11105-014-0700-x

33. Zhang J, Xue B, Gai M, Song S, Jia N, Sun H. Small RNA and Transcriptome Sequencing Reveal a Potential miRNA-Mediated Interaction Network That Functions during Somatic Embryogenesis in Lilium pumilum DC. Fisch. Frontiers in Plant Science. 2017;8: 566-. doi:10.3389/fpls.2017.00566

34. Alejandri-Ramírez ND, Chávez-Hernández EC, Contreras-Guerra JL, Reyes JL, Dinkova TD. Small RNA differential expression and regulation in Tuxpeño maize embryogenic callus induction and establishment. Plant Physiol Biochem. 2018;122: 78–89. doi:10.1016/j.plaphy.2017.11.013

35. Zhou L, Chen J, Li Z, Li X, Hu X, Huang Y, et al. Integrated profiling of microRNAs and mRNAs: microRNAs located on Xq27.3 associate with clear cell renal cell carcinoma. PLoS ONE. 2010;5: e15224. doi:10.1371/journal.pone.0015224

36. Chen C, Ridzon DA, Broomer AJ, Zhou Z, Lee DH, Nguyen JT, et al. Real-time quantification of microRNAs by stem–loop RT–PCR. Nucleic Acids Research. 2005;33: e179. doi:10.1093/nar/gni178

37. Tezuka T, Harada M, Johkan M, Yamasaki S, Oda M. Effects of Auxin and Cytokinin on In Vivo Adventitious Shoot Regeneration from Decapitated Tomato Plants. Hortscience A Publication of the American Society for Horticultural Science. 2011;46: 1661–1665. doi:10.21273/HORTSCI.46.12.1661

38. Plastira VA, Perdikaris AK. EFFECT OF GENOTYPE AND EXPLANT TYPE IN REGENERATION FREQUENCY OF TOMATO IN VITRO. Acta Horticulturae. 1997; 231–234. doi:10.17660/ActaHortic.1997.447.47

39. Sun H-J, Uchii S, Watanabe S, Ezura H. A highly efficient transformation protocol for Micro-Tom, a model cultivar for tomato functional genomics. Plant Cell Physiol. 2006;47: 426–431. doi:10.1093/pcp/pci251

40. Ehlert B, Schöttler MA, Tischendorf G, Ludwig-Müller J, Bock R. The paramutated SULFUREA locus of tomato is involved in auxin biosynthesis. J Exp Bot. 2008;59: 3635–3647. doi:10.1093/jxb/ern213

41. Park J, Lee Y, Martinoia E, Geisler M. Plant hormone transporters: what we know and what we would like to know. Bmc Biology. 2017;15: 93. doi:10.1186/s12915-017-0443-x

42. Hartig K, Beck E. Assessment of lovastatin application as tool in probing cytokinin-mediated cell cycle regulation. Physiologia Plantarum. 2005;125: 260–267. doi:10.1111/j.1399-3054.2005.00556.x

43. Crowell DN, Salaz MS. Inhibition of Growth of Cultured Tobacco Cells at Low Concentrations of Lovastatin Is Reversed by Cytokinin. Plant Physiology. 1992;100: 2090–2095. doi:10.1104/pp.100.4.2090

44. Morris SE, Turnbull CG, Murfet IC, Beveridge CA. Mutational analysis of branching in pea. Evidence that Rms1 and Rms5 regulate the same novel signal. Plant Physiol. 2001;126: 1205–1213. doi:10.1104/pp.126.3.1205

45. Kuroha T, Kato H, Asami T, Yoshida S, Kamada H, Satoh S. A trans-zeatin riboside in root xylem sap negatively regulates adventitious root formation on cucumber hypocotyls. J Exp Bot. 2002;53: 2193–2200. doi:10.1093/jxb/erf077

46. Kudoyarova GR, Vysotskaya LB, Cherkozyanova A, Dodd IC. Effect of partial rootzone drying on the concentration of zeatin-type cytokinins in tomato (Solanum lycopersicum L.) xylem sap and leaves. J Exp Bot. 2007;58: 161–168. doi:10.1093/jxb/erl116

47. Hirose N, Takei K, Kuroha T, Kamada-Nobusada T, Hayashi H, Sakakibara H. Regulation of cytokinin biosynthesis, compartmentalization and translocation. J Exp Bot. 2008;59: 75–83. doi:10.1093/jxb/erm157

48. Takei K, Yamaya T, Sakakibara H. Arabidopsis CYP735A1 and CYP735A2 encode cytokinin hydroxylases that catalyze the biosynthesis of trans-Zeatin. J Biol Chem. 2004;279: 41866–41872. doi:10.1074/jbc.M406337200

49. Williams L, Grigg SP, Xie M, Christensen S, Fletcher JC. Regulation of Arabidopsis shoot apical meristem and lateral organ formation by microRNA miR166g and its AtHD-ZIP target genes. Development. 2005;132: 3657–3668. doi:10.1242/dev.01942

50. Boualem A, Laporte P, Jovanovic M, Laffont C, Plet J, Combier J-P, et al. MicroRNA166 controls root and nodule development in Medicago truncatula. Plant J. 2008;54: 876–887. doi:10.1111/j.1365-313X.2008.03448.x

51. Li Z-X, Li S-G, Zhang L-F, Han S, Li W-F, Xu H, et al. Over-expression of miR166a inhibits cotyledon formation in somatic embryos and promotes lateral root development in seedlings of Larix leptolepis. Plant Cell, Tissue and Organ Culture (PCTOC). 2016;127: 1–13. doi:10.1007/s11240-016-1071-9

52. Jones-Rhoades MW, Bartel DP. Computational Identification of Plant MicroRNAs and Their Targets, Including a Stress-Induced miRNA. Molecular Cell. 2004;14: 787–799. doi:10.1016/j.molcel.2004.05.027

53. Oda Y, Mimura T, Hasezawa S. Regulation of secondary cell wall development by cortical microtubules during tracheary element differentiation in Arabidopsis cell suspensions. Plant Physiol. 2005;137: 1027–1036. doi:10.1104/pp.104.052613

54. Nagasaki H, Jun-ichi Ltoh, Hayashi K, Hibara KI, Satoh-Nagasawa N, Nosaka M, et al. The small interfering RNA production pathway is required for shoot meristem initiation in rice. Proceedings of the National Academy of Sciences of the United States of America. 2007;104: p.14867–14871. doi:10.1073/pnas.0704339104

55. Dinkova TD. Maize miRNA and target regulation in response to hormone depletion and light exposure during somatic embryogenesis. Frontiers in Plant Science. 2015;6: 14.

56. Chu Z, Chen Junying, Xu Haixia, Dong Zhongdong, Chen Feng, Cui Dangqun. Identification and Comparative Analysis of microRNA in Wheat (Triticum aestivum L.) Callus Derived from Mature and Immature Embryos during In vitro Culture. Frontiers in Plant Science. 2016;7. doi:10.3389/fpls.2016.01302

57. Ge F, Luo X, Huang X, Zhang Y, He X, Liu M, et al. Genome-wide analysis of transcription factors involved in maize embryonic callus formation. Physiol Plantarum. 2016;158: 452–462. doi:10.1111/ppl.12470

58. Prakash AP, Kumar PP. PkMADS1 is a novel MADS box gene regulating adventitious shoot induction and vegetative shoot development in Paulownia kawakamii. Plant Journal. 2002;29: 141–151. doi:10.1046/j.0960-7412.2001.01206.x

59. Tanaka N, Itoh H, Sentoku N, Kojima M, Sakakibara H, Izawa T, et al. The COP1 Ortholog PPS Regulates the Juvenile-Adult and Vegetative-Reproductive Phase Changes in Rice. The Plant Cell. 2011;23: 2143–2154.

60. Tian F, Li X, Wu Y, Xia K, Jie O, Zhang M, et al. Rice osa-miR171c Mediates Phase Change from Vegetative to Reproductive Development and Shoot Apical Meristem Maintenance by Repressing Four OsHAM Transcription Factors. Plos One. 2015;10: e0125833-. doi:10.1371/journal.pone.0125833

61. Yang Q, Liu S, Han X, Ma J, Deng W, Wang X, et al. Integrated transcriptome and miRNA analysis uncovers molecular regulators of aerial stem-to-rhizome transition in the medical herb Gynostemma pentaphyllum. BMC Genomics. 2019;20: 865. doi:10.1186/s12864-019-6250-8

62. Ma J, Zhao P, Liu S, Yang Q, Guo H. The Control of Developmental Phase Transitions by microRNAs and Their Targets in Seed Plants. Int J Mol Sci. 2020;21. doi:10.3390/ijms21061971

63. Howell SH, Lall S, Che P. Cytokinins and shoot development. Trends in Plant Science. 2003;8: 0–459. doi:10.1016/s1360-1385(03)00191-2

64. Kieber JJ, Ferreira FJ. Cytokinin signaling. Current Opinion in Plant Biology. 2005;8: 518–525. doi:10.1016/j.pbi.2005.07.013

65. Hwang I, Sheen J, Müller B. Cytokinin signaling networks. Annu Rev Plant Biol. 2012;63: 353–380. doi:10.1146/annurev-arplant-042811-105503

66. Siddiqui ZH, Abbas ZK, Ansari MW, Khan MN. The role of miRNA in somatic embryogenesis. Genomics. 2019;111: 1026–1033. doi:10.1016/j.ygeno.2018.11.022

67. Madhuri Gandikota, Rainer P. Birkenbihl, Susanne Höhmann. The miRNA156/157 recognition element in the 3¢ UTR of the Arabidopsis SBP box gene SPL3 prevents early flowering by translational inhibition in seedlings. Plant Journal. 2007 [cited 3 Apr 2020]. doi:10.1111/j.1365-313X.2006.02983.x

68. Cardon GH, Höhmann S, Nettesheim K, Saedler H, Huijser P. Functional analysis of the Arabidopsis thaliana SBP-box gene SPL3: A novel gene involved in the floral transition. Plant Journal. 1997;12: 367–377. doi:10.1046/j.1365-313X.1997.12020367.x

69. Wu G, Poethig RS. Temporal regulation of shoot development in Arabidopsis thaliana by miR156 and its target SPL3. Development. 2006;133: 3539–3547. doi:10.1242/dev.02521

70. Wang JW, Schwab R, Czech B, Mica E, Weigel D. Dual Effects of miR156-Targeted SPL Genes and CYP78A5/KLUH on Plastochron Length and Organ Size in Arabidopsis thaliana. Plant Cell. 2008;20: 1231–1243. doi:10.1105/tpc.108.058180

71. Unte U, Sorensen A-M, Pesaresi P, Gandikota M, Leister D, Saedler H, et al. SPL8, an SBP-Box Gene That Affects Pollen Sac Development in Arabidopsis. The Plant cell. 2003;15: 1009–19. doi:10.1105/tpc.010678

72. Zhang Y, Schwarz S, Saedler H, Huijser P. SPL8, a local regulator in a subset of gibberellin-mediated developmental processes in Arabidopsis. Plant Molecular Biology. 2007;63: 429–439. doi:10.1007/s11103-006-9099-6

73. Chuck G, Whipple C, Jackson D, Hake S. The maize SBP-box transcription factor encoded by tasselsheath4 regulates bract development and the establishment of meristem boundaries. Development. 2010;137: 1243–1250. doi:10.1242/dev.048348

74. Rhoades MW, Reinhart BJ, Lim LP, Burge CB, Bartel B, Bartel DP. Prediction of Plant MicroRNA Targets. Cell. 2002;110: 0–520. doi:10.1016/s0092-8674(02)00863-2

75. Liu M, Wu X, Long J, Guo W-W. Genomic characterization of miR156 and SQUAMOSA promoter binding protein-like genes in sweet orange (Citrus sinensis). Plant Cell, Tissue and Organ Culture (PCTOC). 2017;130: 1–14. doi:10.1007/s11240-017-1207-6

76. Zhang T-Q, Lian H, Tang H, Dolezal K, Zhou C-M, Yu S, et al. An intrinsic microRNA timer regulates progressive decline in shoot regenerative capacity in plants. Plant Cell. 2015;27: 349–360. doi:10.1105/tpc.114.135186

77. Gao Y, Li D, Zhang L-L, Borthakur D, Li Q-S, Ye J-H, et al. MicroRNAs and their targeted genes associated with phase changes of stem explants during tissue culture of tea plant. Scientific Reports. 2019;9: 20239. doi:10.1038/s41598-019-56686-3

78. Jung J-H, Park C-M. MIR166/165 genes exhibit dynamic expression patterns in regulating shoot apical meristem and floral development in Arabidopsis. Planta. 2007;225: 1327–1338. doi:10.1007/s00425-006-0439-1

79. Wong CE, Zhao Ying-Tao, Wang Xiu-Jie, Larry C, Wang Zhong-Hua, Farzad H, et al. MicroRNAs in the shoot apical meristem of soybean. Journal of Experimental Botany. 2011; 8. doi:10.1093/jxb/erq437

80. Turchi L, Baima S, Morelli G, Ruberti I. Interplay of HD-Zip II and III transcription factors in auxin-regulated plant development. J Exp Bot. 2015;66: 5043– 5053. doi:10.1093/jxb/erv174

81. Brandt R, Salla-Martret M, Bou-Torrent J, Musielak T, Stahl M, Lanz C, et al. Genome-wide binding-site analysis of REVOLUTA reveals a link between leaf patterning and light-mediated growth responses. Plant J. 2012;72: 31–42. doi:10.1111/j.1365-313X.2012.05049.x

82. Zhong R, Ye ZH. Alteration of auxin polar transport in the Arabidopsis ifl1 mutants. Plant Physiol. 2001;126: 549–563. doi:10.1104/pp.126.2.549

83. Huang T, Harrar Y, Lin C, Reinhart B, Newell NR, Talavera-Rauh F, et al. Arabidopsis KANADI1 acts as a transcriptional repressor by interacting with a specific cis-element and regulates auxin biosynthesis, transport, and signaling in opposition to HD-ZIPIII factors. Plant Cell. 2014;26: 246–262. doi:10.1105/tpc.113.111526

84. Li ZX, Zhang LF, Li WF, Qi LW, Han SY. MIR166a Affects the Germination of Somatic Embryos in Larixleptolepis by Modulating IAA Biosynthesis and Signaling Genes. Journal of Plant Growth Regulation. 2017 [cited 1 May 2020]. doi:10.1007/s00344-017-9693-7

85. Li Z-X, Fan Y-R, Dang S-F, Li W-F, Qi L-W, Han S-Y. LaMIR166a-mediated auxin biosynthesis and signalling affect somatic embryogenesis in Larix leptolepis. Mol Genet Genomics. 2018;293: 1355–1363. doi:10.1007/s00438-018-1465-y

86. Vanneste S, Friml J. Auxin: a trigger for change in plant development. Cell. 2009;136: 1005–1016. doi:10.1016/j.cell.2009.03.001

87. Navarro L, Dunoyer P, Jay F, Arnold B, Dharmasiri N, Estelle M, et al. A plant miRNA contributes to antibacterial resistance by repressing auxin signaling. Science. 2006;312: 436–439. doi:10.1126/science.1126088

88. Si-Ammour A, Windels D, Arn-Bouldoires E, Kutter C, Ailhas J, Meins F, et al. miR393 and secondary siRNAs regulate expression of the TIR1/AFB2 auxin receptor clade and auxin-related development of Arabidopsis leaves. Plant Physiol. 2011;157: 683–691. doi:10.1104/pp.111.180083

89. Wójcik AM, Gaj MD. miR393 contributes to the embryogenic transition induced in vitro in Arabidopsis via the modification of the tissue sensitivity to auxin treatment. Planta. 2016;244: 231–243. doi:10.1007/s00425-016-2505-7

90. Wu M-F, Tian Q, Reed JW. Arabidopsis microRNA167 controls patterns of ARF6 and ARF8 expression, and regulates both female and male reproduction. Development. 2006;133: 4211–4218. doi:10.1242/dev.02602

91. Liu Z, Li J, Wang L, Li Q, Lu Q, Yu Y, et al. Repression of callus initiation by the miRNA-directed interaction of auxin-cytokinin in Arabidopsis thaliana. Plant Journal. 2016 [cited 15 Mar 2020]. doi:10.1111/tpj.13211

92. Sakai H, Honma T, Aoyama T, Sato S, Kato T, Tabata S, et al. ARR1, a transcription factor for genes immediately responsive to cytokinins. Science. 2001;294: 1519–1521. doi:10.1126/science.1065201

93. Buechel S, Leibfried A, To JPC, Zhao Z, Andersen SU, Kieber JJ, et al. Role of A-type ARABIDOPSIS RESPONSE REGULATORS in meristem maintenance and regeneration. Eur J Cell Biol. 2010;89: 279–284. doi:10.1016/j.ejcb.2009.11.016

94. Gordon SP, Heisler MG, Reddy GV, Ohno C, Das P, Meyerowitz EM. Pattern formation during de novo assembly of the Arabidopsis shoot meristem. Development. 2007;134: 3539–3548. doi:10.1242/dev.010298

95. Sunkar R, Zhu J-K. Novel and stress-regulated microRNAs and other small RNAs from Arabidopsis. Plant Cell 16, 2001−2019. Plant Cell. 2004;16: 2001–2019. doi:10.1105/tpc.104.022830

96. Constabel CP, Yip L, Patton JJ, Christopher ME. Polyphenol oxidase from hybrid poplar. Cloning and expression in response to wounding and herbivory. Plant Physiol. 2000;124: 285–295. doi:10.1104/pp.124.1.285

97. Lu S, Li Q, Wei H, Chang M-J, Tunlaya-Anukit S, Kim H, et al. Ptr-miR397a is a negative regulator of laccase genes affecting lignin content in Populus trichocarpa. Proc Natl Acad Sci USA. 2013;110: 10848–10853. doi:10.1073/pnas.1308936110

98. Dauwe R, Morreel K, Goeminne G, Gielen B, Rohde A, Van Beeumen J, et al. Molecular phenotyping of lignin-modified tobacco reveals associated changes in cell-wall metabolism, primary metabolism, stress metabolism and photorespiration. Plant J. 2007;52: 263–285. doi:10.1111/j.1365-313X.2007.03233.x

99. Motose H, Fukuda H, Sugiyama M. Involvement of local intercellular communication in the differentiation of zinnia mesophyll cells into tracheary elements. Planta. 2001;213: 121–131. doi:10.1007/s004250000482

